# Visuospatial n-back test for early detection of mild cognitive impairment in patients with Parkinson’s Disease : an fMRI study

**DOI:** 10.1101/2020.03.03.974725

**Authors:** Shoji Kawashima, Yoko Shimizu, Yoshino Ueki, Noriyuki Matsukawa

## Abstract

**Background:** Cognitive impairment is a common symptom in the patients with Parkinson’s disease (PD). In delineating a therapeutic plan, the early diagnosis of mild cognitive impairment in PD (PD-MCI) is important. Patients with PD-MCI have severe impairment in frontal executive function and/or visuospatial recognition. However, the clinical assessment of these functions is not routinely performed.

**Method:** In this study, we aimed to clarify the advantage of visuospatial version of the n-back test as a tool for the early detection of neuropsychological change in the patients with PD-MCI. The score of 0-back test reflects visuospatial recognition, and the scores of 1-back and 2-back reflect visuospatial working memory. PD-MCI was classified according to the criteria provided by the Movement Disorder Society Task Force for mild cognitive impairment in PD. We recruited 13 patients with PD-MCI, and 15 patients with cognitive normal PD. Using functional MRI (fMRI), we also aimed to clarify the specific brain regions associated with the impairment of visuospatial working memory.

**Result:** We demonstrated that the correct answer rate of patients with PD-MCI was lower in the 2-back test than patients with PD-CN. However, we did not find statistical difference in the 0-back test. These results indicate the preservation of visuospatial recognition and the impairment of visuospatial working memory in the patients with PD-MCI. We revealed the reduced activation within the middle frontal gyrus (MFG) and the inferior parietal lobule (IPL) during the 2-back test in the patients with PD-MCI. It may be associated with the severity of cortico-striatal dysfunction in the dopaminergic neural network which is associated with Lewy body pathology.

**Conclusion:** The visuospatial n-back test has advantages for use in rapid and early detection of impaired visual recognition and working memory. The combination of functional neuroimaging and neuropsychological tests may provide markers for the increased risk of dementia before the development of an irreversible disease-specific pathology.

## Introduction

Striatal dopamine depletion due to the degeneration of nigrostriatal dopaminergic neurons causes motor disturbances in the patients with Parkinson’s disease (PD). The cognitive impairment is common symptoms of PD as well as motor disturbances. The movement disease society (MDS) has published the clinical criteria for dementia in PD (PDD) in 2007 [1], and the MDS Task Force has provided the criteria for mild cognitive impairment (MCI) in PD (PD-MCI) in 2012 [2]. MCI is a transitional stage between normal ageing and dementia that has been used to detect and treat early dementia [3]. In a recent cohort study, 20% of *de novo* patients with PD were classified as PD-MCI at baseline, and 39% of patients with baseline or incident PD-MCI progressed to dementia during the 5-year follow-up period [4]. Therefore, the early diagnosis of PD-MCI is important for clinical management.

The Montreal Cognitive Assessment (MoCA) is more sensitive than MMSE in testing global cognition in the patients with PD [5]. The previous studies have revealed that impairments in frontal executive function and visual recognition are associated with dementia in PD [6, 7]. Working memory, which is responsible for the short-term storage and online manipulation of information necessary for higher cognitive function, is important; impaired working memory may result in disrupted activities of daily living. The trail-making or Stroop tests are commonly used to assess executive function; however, clinical assessments of working memory were not routinely performed.

The verbal n-back test has been used in neuroimaging studies that incorporate functional MRI (fMRI) to explore brain activation associated with working memory processing [8]. The visuospatial version of the n-back test was used in fMRI studies for normal subjects [9, 10]. This test assesses visuospatial function and working memory, which may reflect the early pathological progression to PDD.

In this study, using fMRI, we aimed to investigate neuroimaging characteristics associated with visuospatial working memory in PD-MCI, compared with cognitively normal PD. We hypothesised that the patients with PD-MCI show decreased brain activation in association with impaired visuospatial working memory.

## Methods

### Subjects

This study was approved by the Institutional Review Board of Nagoya City University Hospitals. We enrolled 28 right-handed patients with PD (18 males, 10 females) and 12 age-matched normal subjects in the study. All the patients included in the study fulfilled the UK PD Society Brain Bank Criteria for clinical diagnosis, and none had a disease other than PD that affected motor and cognitive function. The patients were assessed with the Unified Parkinson’s Disease Rating Scale (UPDRS) and the Hoehn and Yahr Scale at the time of inclusion in the study. The patients were excluded if they had dementia according to the criteria for PD dementia provided by the Movement Disorder Society Task Force [1]. We excluded also patients if they had depression, severe insomnia, severe hearing loss, or any other disease that might severely interfere with the fMRI. All the subjects provided written informed consent prior to the data acquisition.

PD-MCI was classified according to the criteria provided by the Movement Disorder Society Task Force for mild cognitive impairment in PD. In brief, PD-MCI was diagnosed when patients’ scores were 1.5 SD below the normative mean score in 2 cognitive domains on at least one impaired test in two different cognitive domains [2]. The patients were classified as PD-CN when scores were within 1.5 SD of the normative mean score.

### Neuropsychological test

Movement disorder specialists performed the complete neuropsychological battery in all the patients and the normal subjects. The MMSE and MoCA were used to assess global cognitive impairment. Psychomotor speed and attention of subjects were tested with Trail-Making Test Part A (TMT-A) and the Paced Auditory Serial Addition Test (PASAT). Attention and rapid set shifting were tested with Trail-Making Test Part B (TMT-B). Verbal memory was tested with the Auditory Verbal Learning Test (AVLT); language was tested with the Verbal Fluency Test, and visuospatial function was tested with the clock-copying test and the visuospatial version of the 0-back test. Depressive mental status was scored using the Beck Depression Inventory (BDI).

### fMRI protocol

The patients were asked to perform the visuospatial n-back task with 3 load levels during the fMRI. The stimuli were white squares randomly presented in 1 of 8 spatial locations on a screen, through a mirror positioned on a head-coil. The presentation of the stimuli was controlled by a program (Presentation software) that initiated the acquisition of the MRI and the behavioural data. For the 0-back test, the subjects were instructed to press the left button with their index finger when a white square was presented. For the 1-back test, the patients pressed the left button whenever a stimulus was presented in the same location as the previous stimulus. For the 2-back test, when the stimulus was presented in any other location, the patients were instructed to press the right button with the middle finger. The patients were instructed to press the right button when the stimulus appeared in any other location.

To detect brain regions activated in the subjects performing the visuospatial n-back task, we used the block-design fMRI, which alternated between n-back test and rest conditions. For n-back task conditions, the white square was randomly presented for 2 s in 1 of 8 possible locations on screen; a black screen was presented for 1 s after stimulus presentation. Each test condition consisted of 15 trials over the course of 45 s; each rest condition lasted for 15. During a single scan, each test condition was repeated 4 times, in numerical order (0-1-2 back). Thus, each condition included 60 trials (Fig. 1).

**Fig 1.**
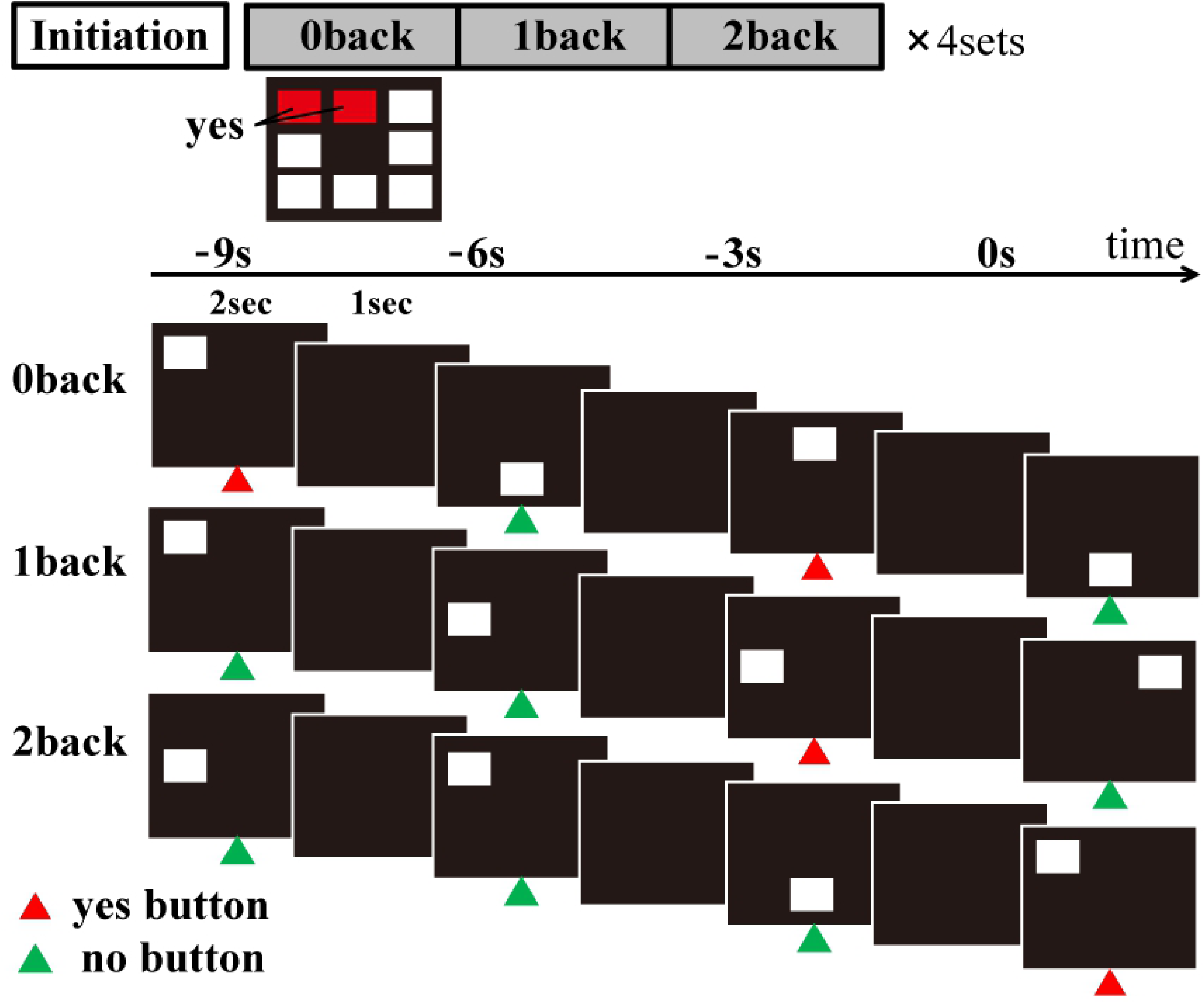
Study protocol. The figure shows the protocol of n-back task used in this study. To detect the activated brain regions associated with the visuospatial n-back task, we used a blocked design fMRI, alternating n-back test conditions and rest conditions. In n-back task conditions, the white square was presented for 2 s at random in 1 of 8 possible locations on screen, and black screen was presented for 1 s after the presentation of stimuli. In the 0-back test, the subjects were instructed to press the left button with their index finger when the white square was presented in predetermined locations. In the 1-back test, the subjects were instructed to press the left button when the stimulus presented in the same location as the previous one. In the 2-back test, the subjects were instructed to press the right button with middle finger when the stimulus presented in any other location. When in any other location, the subjects were instructed to press right button.

### Image data acquisition and analysis

All MRI were acquired with a Siemens Skyla syngo MR E11 3.0 T scanner (Siemens, Germany). High-resolution T1-weighted images were acquired via volumetric 3D spoiled gradient recall sequence. Acquisition parameters were as follows: repetition time (TR) = 1900 ms, echo time (TE) = 2.43 ms, flip angle (FA) = 9, field of view (FOV) = 256 × 256 mm, slice thickness = 1 mm, slice gap = 0, voxel size = 1 × 1 × 1 mm, number of slices = 176. The fMRI measurements were performed using a gradient echo EPI sequence: repetition time (TR) = 2500 ms, echo time (TE) = 30 ms, slice thickness = 3 mm, with matrix size of 64 × 64 and field of view of 192 × 192 mm, resulting in voxel size of 3 × 3 × 3 mm.

All the images were pre-processed and analysed with Matlab (version X, The Mathworks Inc, Natick, MA) and SPM8 software (Department of Cognitive Neurology, London). Images were realigned to correct for movement and normalised to Montreal Neurologic Institute (MNI) space. The transformed image data were smoothed with a Gaussian philtre (full width at half-maximum = 10 mm).

The image data were analysed with a random effects procedure and a parametric model to identify the brain areas where the activation correlated with the task. Group comparisons between PD-MCI and PD-CN were performed for 3 loads of the n-back test, the 1-back vs. 0-back condition and the 2-back vs. 0-back condition. The correlations between score on the n-back test and task-related activation were analysed for all the patients. The statistical significance was established at P < 0.01 (uncorrected) with cluster size >50 voxels in group analysis, and P < 0.001 (uncorrected) with cluster size >50 in correlation analysis.

### Statistical analysis

The group differences in the clinical profiles were analysed by the t-test as appropriate. The scores on n-back tests and reaction times were analysed using one-way analysis of variance (ANOVA), because each load level required several neuropsychological cognitive domains. SPSS version 15.0 was used for these analyses. P<0.05 was considered significant.

## Results

### Clinical profiles of patients with PD-MCI and PD-CN

The demographic and clinical profiles of patients with PD are summarised in Table 1. A total of 13 patients were classified as MCI (PD-MCI); 15 patients were classified as cognitively normal (PD-CN). The UPDRS motor scores of the patients with PD-MCI were significantly higher than those of the patients with PD-CN. Age, sex, disease duration, Hoehn and Yahr stage and BDI were not different between groups. The patients in both groups received similar daily doses of LDOPA, total LDOPA and L-dopa equivalent daily dose of dopamine agonist (LEDD). In the comparisons of global cognition, the total MMSE scores were similar in both of the groups; however, the MoCA scores were significantly lower in the patients with PD-MCI, compared to the patients with PD-CN. In comparisons of neuropsychological subdomains, scores on the PASAT, AVLT and Verbal Fluency Test were lower in the patients with PD-MCI, compared to the patients with PD-CN. Patients with PD-MCI took significantly longer to complete the TMT-A and TMT-B. There was no difference in the scores on pentagon copy test, used for screening of visuospatial impairment.

**Table 1.** Baseline characteristics of the patients with PD-MCI and PD-CN

|  | **PD-MCI** | **PD-CN** |  |
| --- | --- | --- | --- |
|  | **n = 13** | **n = 15** | **P** |
| Age | 69.4 ± 2.8 | 67.3 ± 4.6 | N.S. |
| Men | 8 | 10 | N.S. |
| Disease Duration (year) | 5.0 ± 3.2 | 5.4 ± 3.5 | N.S. |
| HY stage | 2.1 ± 0.8 | 2.1 ± 0.7 | N.S. |
| UPDRS part3 | 19.8 ± 8.6 | 10.1 ± 5.2 | p < 0.01 |
| LDOPA (mg) | 262 ± 153 | 223 ± 90 | N.S. |
| LEDD (mg) | 296 ± 199 | 287 ± 151 | N.S. |
| MMSE | 26.9 ± 1.8 | 28.1 ± 1.9 | N.S. |
| MoCA | 21.1 ± 3.0 | 24.3 ± 2.8 | p < 0.01 |
| TMT-A (second) | 65.6 ± 21.0 | 43.8 ± 11.2 | p < 0.01 |
| TMT-B (second) | 276 ± 107.5 | 111 ± 50.4 | p < 0.001 |
| PASAT | 21.2 ± 6.1 | 27.2 ± 6.2 | p < 0.05 |
| AVLT | 5.5 ± 3.2 | 9.1 ± 2.9 | p < 0.01 |
| Verbal fluency | 12 ± 2.9 | 15.5 ± 3.1 | p < 0.01 |
| Copy of pentagon | $0.92 \pm 0.28$ | $1.0 \pm 0$ | N.S. |
| BDI | $12.1 \pm 7.6$ | $9.1 \pm 7.0$ | N.S. |

Duration: Disease duration; HY stage: Hoehn and Yahr stage; UPDRS motor: motor sections of united PD rating scale; LEDD: L-dopa equivalent daily dose of dopamine agonist. Calculation of LDDD for each patient was based on theoretical equivalence to L-dopa as follows: L-dopa dose + L-dopa dose × 1/3 [if on entacapone + bromocriptine (mg) × 10 + cabergoline or pramipexole (mg) × 67 + ropinirole (mg) × 20 + pergolide (mg) × 100 + apomorphine (mg) × 8]. MoCA, Montreal Cognitive Assessment, PASAT, Paced Auditory Serial Addition Test; AVLT, Auditory Visual Learning Test; BDI, Beck Depression Inventory; N.S., not significant

### Comparisons of performance on visuospatial n-back test

Correct answer rates and reaction times for all 3 task conditions are summarised in Fig 2. ANOVA revealed a significant difference between groups for scores on the 2-back test. PD-MCI patients scored lower than PD-CN patients (F(26)=6.001, P=0.021). Scores on the 0-back and 1-back tests were similar between the groups (0-back: P=0.11; 1-back: P=0.07).

**Fig 2.**
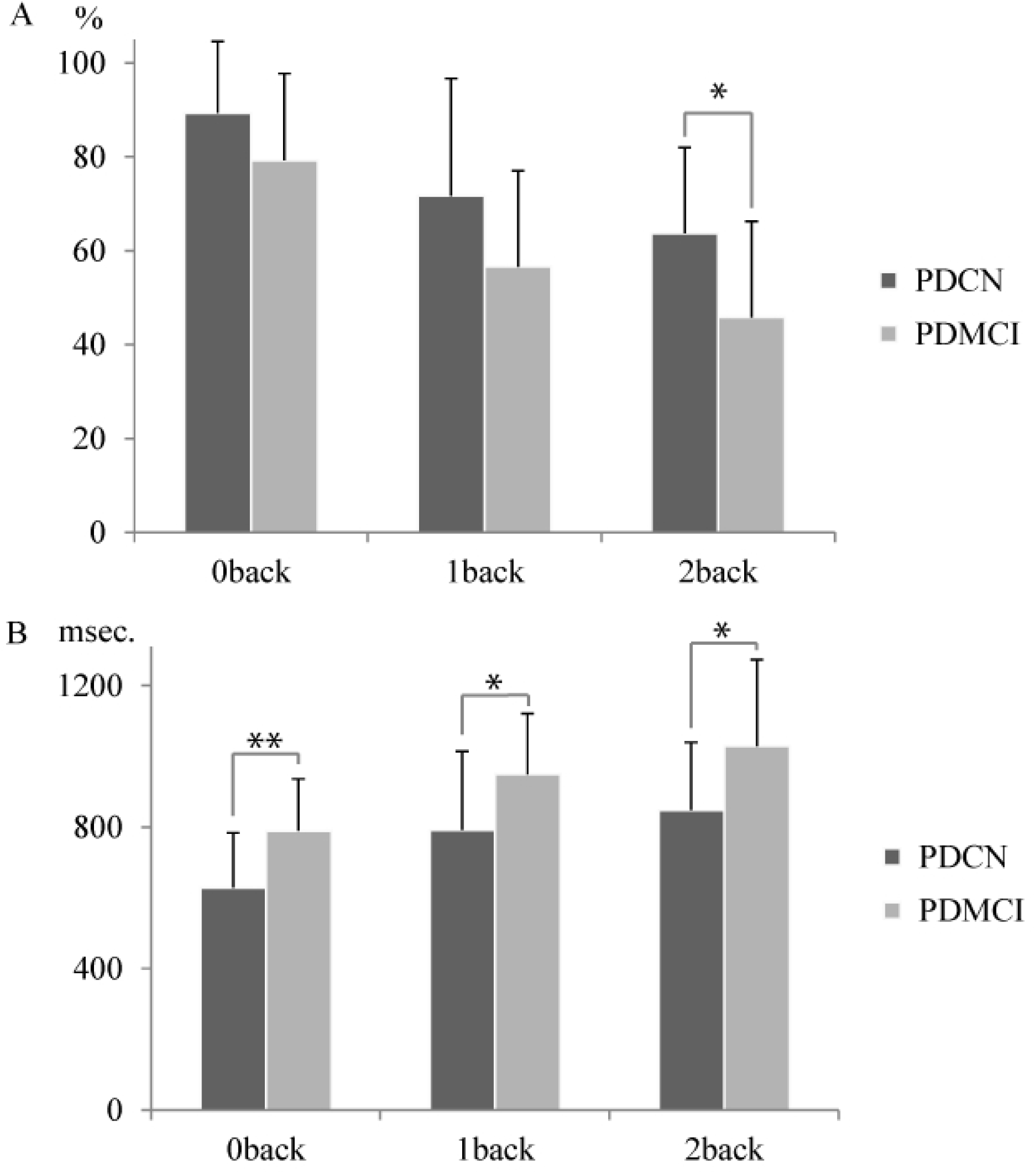
Performance of the visuospatial n-back test. A. The figure shows rates of correct answers for the patients with PD-CN and PD-MCI. The scores on the 2-back test of PD-MCI were significantly lower than that of PD-CN. B. The figure shows the response time to press correct button. The patients with PD-MCI took significantly longer time to respond. * P<0.05, ** P<0.01

The reaction times on the 0-back test were longer in the PD-MCI group, compared with the PD-CN group (F[25]=5.469, P<0.05). The reaction times were also longer for the 1-back and 2-back test in the PD-MCI group, compared with the PD-CN group.

### Correlation between scores on the n-back test and other factors

The correlations between scores on the n-back test and other factors are presented in Table 2. There was a significant negative correlation between the scores on the 1-back test and the score on UPDRS Part 3 (R=-0.41, P<0.05) and between the scores on the 2-back test and the score on UPDRS Part 3 (R=-0.42, P<0.05). Additionally, we found positive correlations between scores on MoCA and the 1-back test (R=0.39, P<0.05), as well as between scores on MoCA and the 2-back test (R=0.41, P<0.05). We also found a negative correlation between scores on the TMT-B and the 0-back test (R=-0.50, P<0.01), however, there was no correlation between scores on TMT-A and any load of the n-back test.

**Table 2.** Correlations between scores on the n-back test and other factors

|  |  | **0-back** | **1-back** | **2-back** | **PASAT** | **TMT-B** |
| --- | --- | --- | --- | --- | --- | --- |
| **MoCA** | Peason' R |  | 0.39 | 0.413 | 0.35 | −0.54 |
|  | P value | N.S. | 0.040 | 0.029 | 0.041 | 0.003 |
| **TMT-A** | Peason' R |  |  |  |  | 0.524 |
|  | P value | N.S. | N.S. | N.S. | N.S. | 0.004 |
| **TMT-B** | Peason' R | −0.495 | −0.5 |  | −0.5 | 1 |
|  | P value | 0.007 | 0.007 | N.S. | 0.007 |  |
| **UPDRS**<br>**part3** | Peason' R |  | −0.406 | −0.415 | −0.406 | 0.685 |
|  | P value | N.S. | 0.032 | 0.028 | 0.032 | 0.001 |

N.S.: not significant

### Functional MRI

The fMRI group analyses were performed to compare the patients with PD-MCI and the patients with PD-CN. Talairach coordinates of the centre of activation and cluster size for significant findings are summarised in Table 3. For the 0-back test, the PD-MCI group presented significantly reduced brain activation within the right inferior frontal gyrus (IFG), left superior frontal gyrus (SFG) and left medial frontal gyrus. For the 1-back test, significant differences between the groups were observed for right IFG, bilateral superior parietal lobule (SPL) and bilateral cuneus. For the 2-back test, the PD-MCI group presented significantly reduced brain activation within right inferior parietal lobule (IPL), right middle frontal gyrus (MFG), left SPL, left lateral globus pallidus, left cerebellar tonsil and left precentral gyrus. For the 2-back versus 0-back condition, the patients with PD-MCI presented reduced activation within the bilateral MFG and IPL, compared to the patients with PD-CN. The images of significant voxels for each load of the n-back test are presented in Fig 3.

**Table 3.**
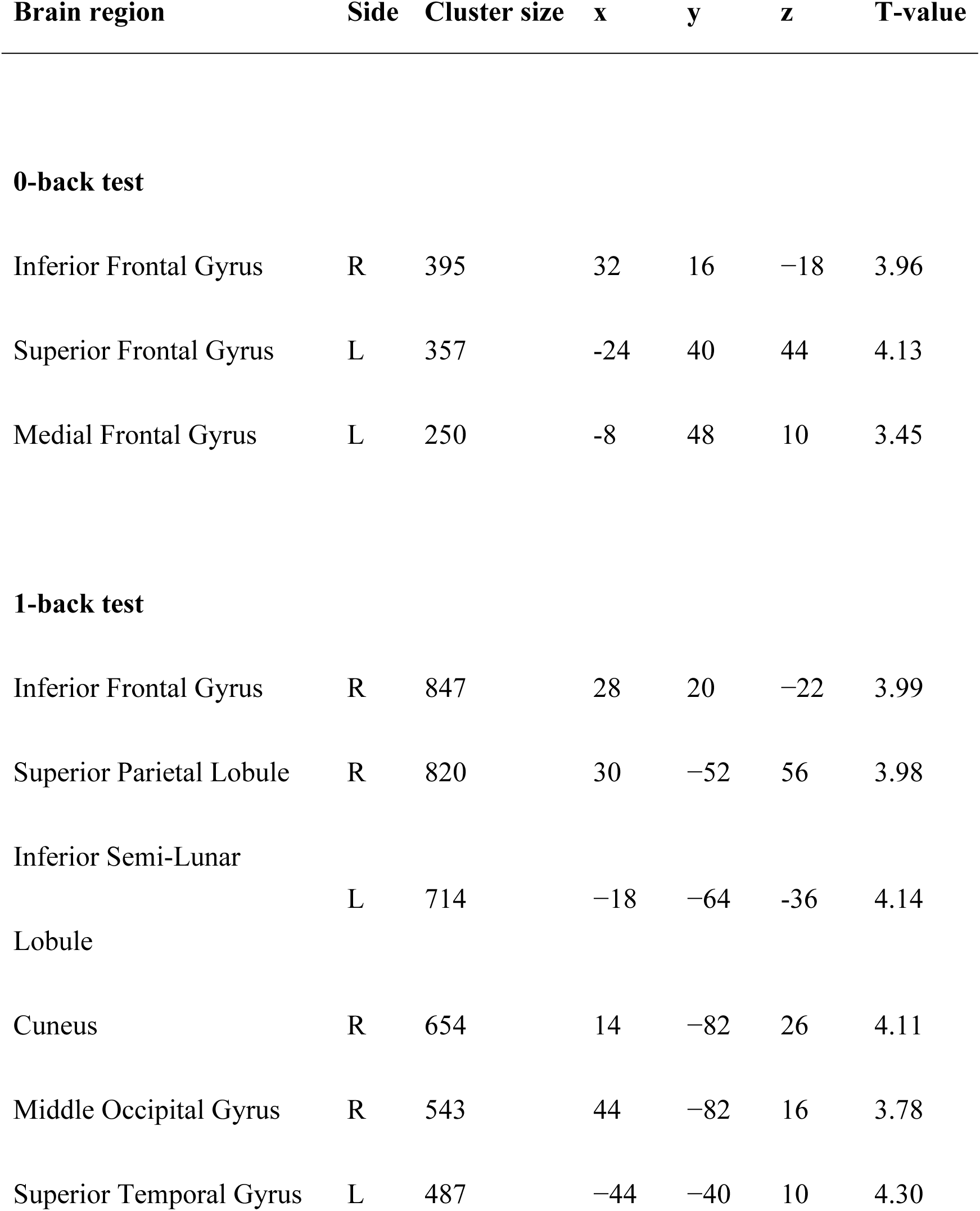
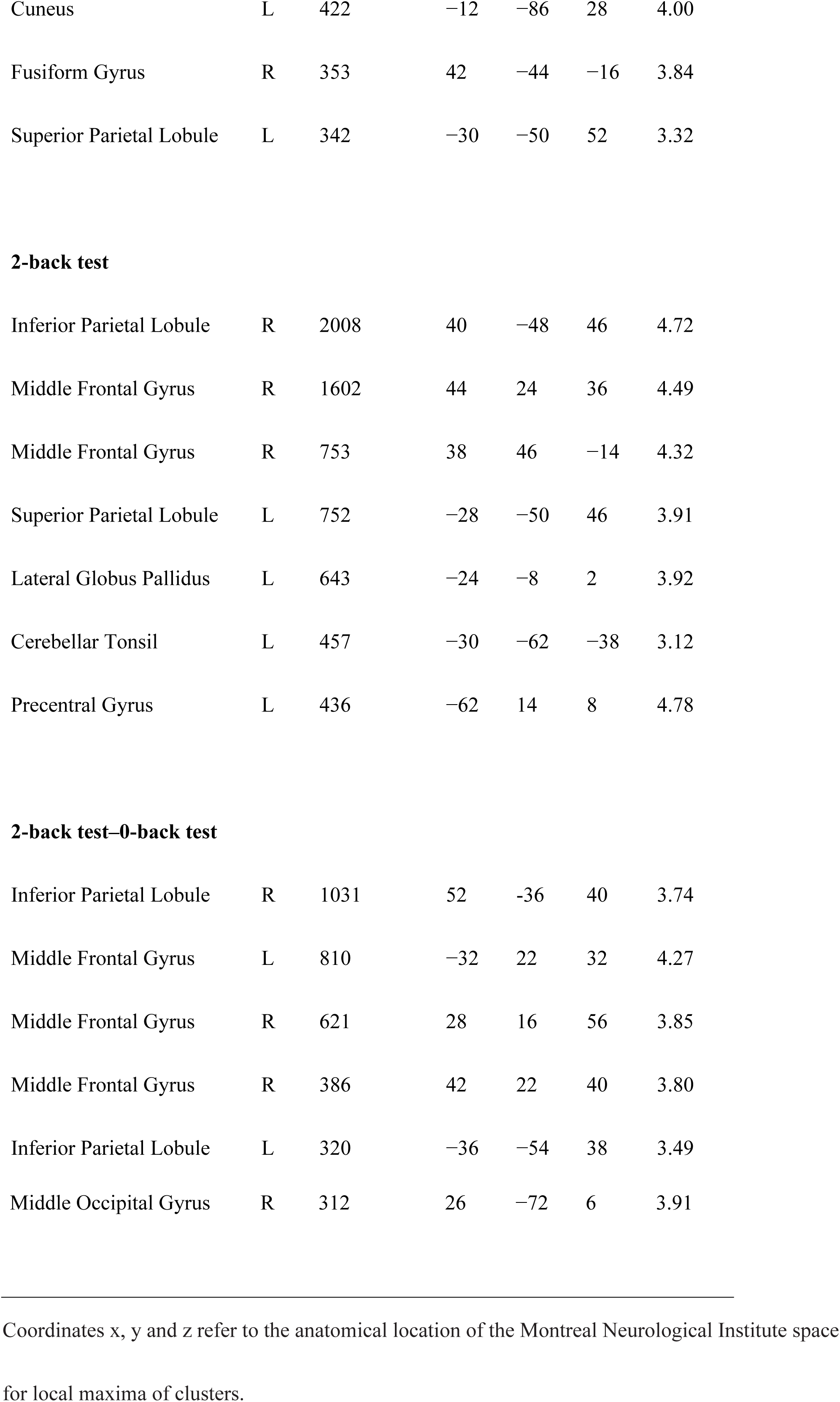
Talairach coordinates of the centre of activation for each load of the n-back test

**Fig 3.**
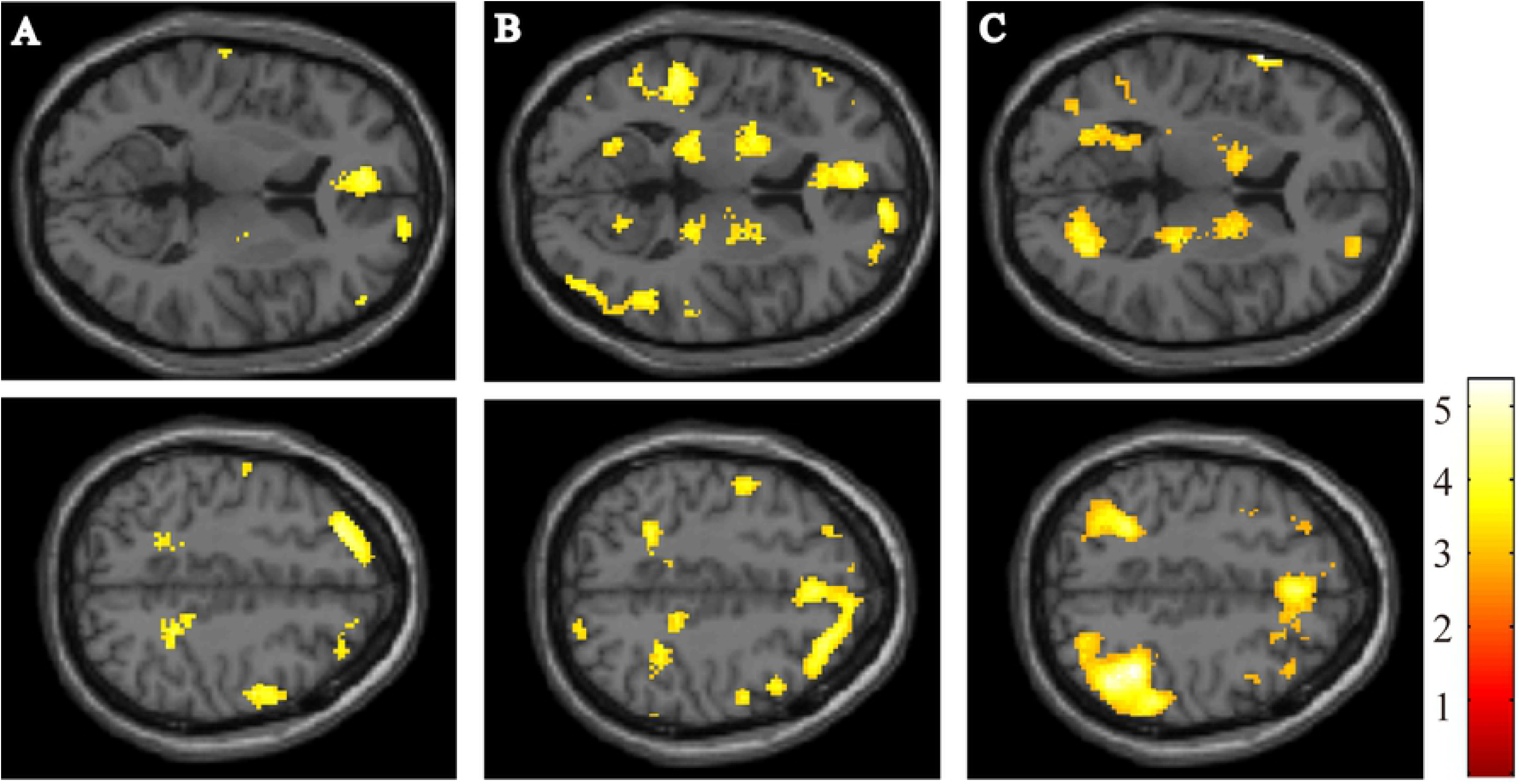
fMRI analyses comparing PD-MCI and PD-CN. The figure shows the results for the 0-back test (A), 1-back test (B) and 2-back test (C). The coloured regions indicate significantly lower brain activation in PD-MCI, as compared with PD-CN. All the images presented at P < 0.01 (uncorrected) with cluster size >50 voxels in analysis.

The correlations between the scores on the n-back test and task-related activation are shown in Fig 4. We found a positive correlation between the score on the 2-back test within the right IPL (Fig 4B) and the scores on the 1-back test within MFG and SPL (Fig 4A).

**Figure 4.**
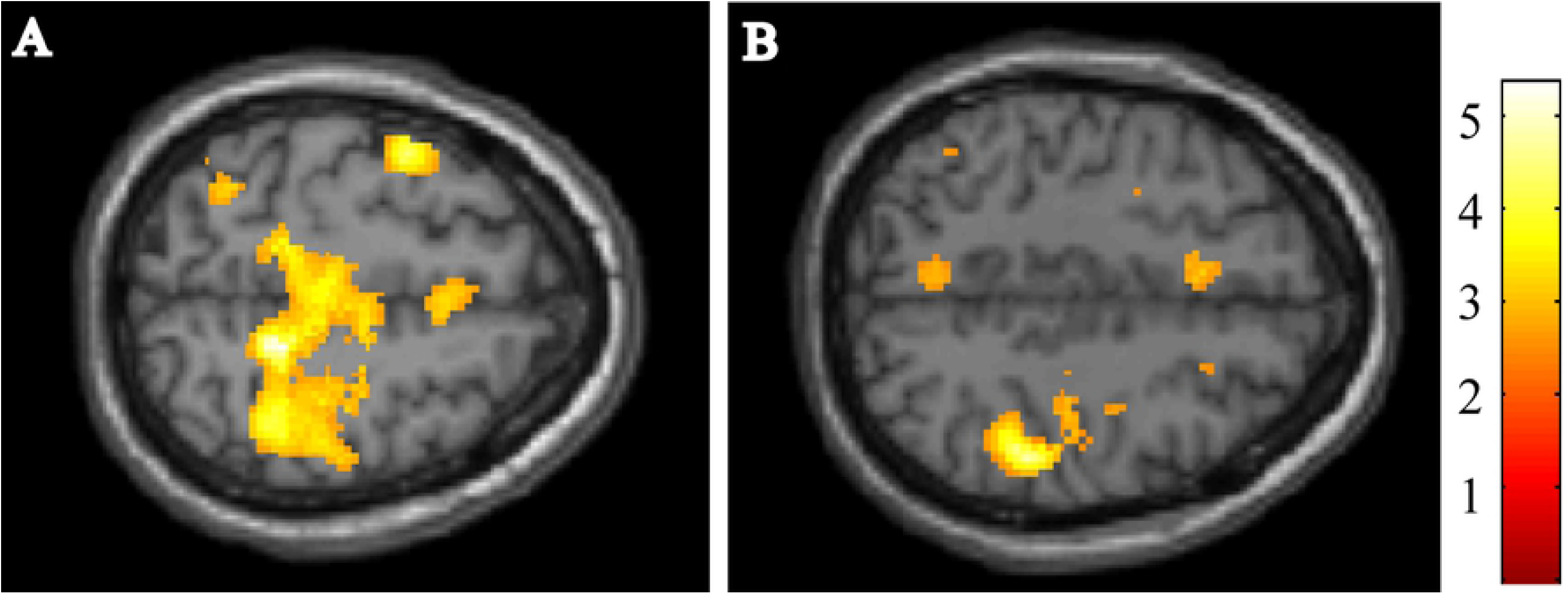
Correlation analyses. A. The image shows the correlation between the scores on the 1-back test and activation during the 1-back test. B. The image shows the correlation between the scores on the 2-back test and activation during the 2-back test.

## Discussion

This study aimed to investigate the impairment of visuospatial working memory in patients with PD-MCI, using a visuospatial version of the n-back test. Memory load may be modulated by maintaining consistency in where visual stimuli are presented. We found that the patients with PD-MCI scored lower on the 2-back test than the patients with PD-CN. However, the scores on the 0-back test did not differ significantly between groups. These results indicate the preservation of visuospatial recognition and impaired visuospatial working memory in PD-MCI.

### Neuropsychological characteristics of PD-MCI

A recent meta-analysis reported that the prevalence of PD-MCI as 26% [11]. Another revealed a prevalence of 55% among patients with mean disease duration > 10 years [12]. Approximately 10% of patients with PD convert from PD-MCI to dementia each year [13]. Another study reported that 59% of patients with persistent PD-MCI at 1 year had converted to dementia during follow-up [4]. This evidence indicates that diagnosis of MCI in patients with PD indicates increased risk for dementia. Therefore, neuropsychological assessment and follow-up are important to establish a therapeutic plan for treatment of PD.

Investigations of all cognitive subdomains require an enormous time commitment from patients and neuropsychologists. MoCA is commonly used for cognitive screening of MCI because it includes all cognitive subdomains, as recommended by the MDS task force [2]. The MoCA test has high sensitivity in detection of executive dysfunction. However, this test has limited accuracy in diagnosis of PD-MCI with formal neuropsychological testing (as advocated in Level 2 criteria provided by the MDS task force) [14].

The diagnosis of PD-MCI is important for clinical management of PD, and the visuospatial n-back task is a valuable marker as a screening of the PD-MCI and network deficits caused by pathological development. In this study, scores on the 1-back and 2-back test correlated positively with MoCA, and scores on the 1-back test correlated negatively with frontal executive function (TMT-B). Therefore, the visuospatial n-back test is suitable for the use of screening for cognitive impairment in PD

### Dysfunction of visuospatial working memory in PD-MCI: roles of MFG and IPL

The development of cognitive impairment in PD is caused by complex pathology. Diffuse Levy bodies [15-17], Alzheimer disease [16, 17], loss of cholinergic neurones [18], loss of medial nigral dopaminergic neurones [19], and serotonergic and noradrenergic deficits [20] are implicated in dementia in PD. However, neuropathological data on PD-MCI remain scarce, and the role of each type of pathology in PD-MCI remains unclear.

A structural MRI study reported that, compared with PD-CN, patients with PD-MCI had more severe cortical thinning in the frontal and temporo-parietal cortices [21]. One study of de novo, newly diagnosed PD, which incorporated brain connectivity analysis of resting-state fMRI, revealed that patients with PD-MCI showed decreased functional connectivity between posterior cingulate cortex and posterior IPL [22].

Here we compared the patients with PD-MCI and the patients with PD-CN. Subtraction of activated regions in the 2-back minus 0-back test showed that functional activation of PD-MCI was reduced within bilateral MFG and IPL, compared with PD-CN. The results presented show the destruction of regions associated with motor performance and visual cognition. Furthermore, the scores on the 2-back test correlated positively with activation in right IPL, and scores on the 1-back test correlated positively with activation in MFG and SPL.

Anatomically, MFG belongs to the dorsolateral prefrontal cortex, which is associated with dopaminergic neurones that connect to the caudate nucleus. A post-mortem autopsy report on patients with PDD demonstrated that patients with PD had reduced levels of dopaminergic transporter in caudate, precuneus and IPL [20]. Reduced activation within MFG and IPL may indicate the severity of cortico-striatal dysfunction in the dopaminergic neural network that develops in association with Levy body pathology. Therefore, the observed regions are specific for impairment of visuospatial working memory in patients with PD-MCI.

### Clinical implications

Working memory impairment is common in patients with PD, even in the early stages. However, various studies report conflicting results of the fMRI during tasks that require working memory. A recent fMRI study that incorporated the n-back task reported that functional activity of de novo patients with PD, compared with controls, was increased in right dorso-lateral prefrontal cortex (including MFG) [23]. The results suggest compensation to maintain behavioural performance in the presence of de novo network deficits. In contrast, Simioni et al. reported that patients with PD who were off dopamine replacement therapy displayed reduced activation in prefrontal and bilateral parietal cortex, compared with controls [24]. The discrepancy in task-related activation may depend on the patient sample and variability in the extent of compensation for network deficits or dysfunction.

Executive dysfunction is associated with increased risk for driving-related accidents among patients with PD. Driving performance is significantly impaired in PD patients, compared with controls [25]. One neuropsychological study showed that working memory (n-back task) and mental flexibility (plus-minus task) are critical for driving safety [26]. In this context, cognitive assessment of executive function, especially visuospatial working memory, may help to detect patients with dementia or PD-MCI having high risk of traffic accident.

### Limitations of the study

This study had some limitations. First, performance of neuropsychological tests for executive function is influenced by several factors. Disease-associated impairment of working memory, as well as psychosis, depression and daytime sleepiness, may affect performance. The visuospatial n-back test, in particular, requires continuous attention. Increased variability in subjects’ performance on the 2-back test, compared with the 1-back test, results in decreased statistical power. Second, although the LDOPA and the LEDD doses were similar between groups, we cannot rule out the possibility that variety of the dopaminergic agonist may have had differing impacts on performance. Third, because of the relatively small sample size, the statistical power for identification of significant voxels in the fMRI analyses was limited.

However, the present study had several strengths. First, visual recognition and working memory were assessed using a single test, with 3 different loads. Second, the results presented here demonstrate reduced activation within MFG and IPL in patients with PD-MCI associated with visuospatial working memory. The observed findings are more regional than those typically presented for normal individuals. The dysfunction of MFG and IPL may predict cognitive impairment in PD.

## Conclusion

The visuospatial n-back test has several advantages for use in rapid and early detection of impaired visual recognition and working memory. This study demonstrated the dysfunction of middle frontal gyrus and inferior parietal lobule in patients with PD-MCI. Combinations of functional neuroimaging and neuropsychological testing are beneficial to identify markers of increased risk for dementia before the development of irreversible disease-specific pathology.

## Acknowledgements

We wish to thank Dr. Satoshi Tanaka for technical support of functional MRI analysis. We are also grateful to the patients who took part in this study.

## Grant

This study was supported in part by Grants-in-Aid for Scientific Research on Priority Areas (Grant number is 15K16362).

## Abbreviations used

fMRI: functional magnetic resonance imaging;
PD: Parkinson disease;
PD-CN: PD with normal cognition;
PD-MCI: PD with mild cognitive impairment

